# Deciphering microbial interactions in synthetic human gut microbiome communities

**DOI:** 10.1101/228395

**Authors:** Ophelia S. Venturelli, Alex C. Carr, Garth Fisher, Ryan H. Hsu, Rebecca Lau, Benjamin P. Bowen, Trent Northen, Adam P. Arkin

## Abstract

The human gut microbiota comprises a dynamic ecological system that contributes significantly to human health and disease. The ecological forces that govern community assembly and stability in the gut microbiota remain unresolved. We developed a generalizable model-guided framework to predict higher-order consortia from time-resolved measurements of lower-order assemblages. This method was employed to decipher microbial interactions in a diverse 12-member human gut microbiome synthetic community. We show that microbial growth parameters and pairwise interactions are the major drivers of multi-species community dynamics, as opposed to context-dependent (conditional) interactions. The inferred microbial interaction network as well as a top-down approach to community assembly pinpointed both ecological driver and responsive species that were significantly modulated by microbial inter-relationships. Our model demonstrated that negative pairwise interactions could generate history-dependent responses of initial species proportions on physiological timescales that frequently does not originate from bistability. The model elucidated a topology for robust coexistence in pairwise assemblages consisting of a negative feedback loop that balances disparities in monospecies fitness levels. Bayesian statistical methods were used to evaluate the constraint of model parameters by the experimental data. Measurements of extracellular metabolites illuminated the metabolic capabilities of monospecies and potential molecular basis for competitive and cooperative interactions in the community. However, these data failed to predict influential organisms shaping community assembly. In sum, these methods defined the ecological roles of key species shaping community assembly and illuminated network design principles of microbial communities.

## INTRODUCTION

Microbes have evolved in diverse microbial communities that occupy nearly every environment on Earth, spanning extreme environments such as acid mine drains and hot springs to multicellular organisms. The gut microbiota is a dense of collection of microorganisms that inhabits the human gastrointestinal tract^1–3^ and performs numerous functions to impact human physiology, nutrition, behavior and development^4–9^. The core functions of the gut microbiota are partitioned among genetically distinct populations and integrated into community-level functions such as resistance to invasion and complex chemical transformations. Such collective functions are realized by the combined interactions of diverse microbial species operating on multiple time and spatial scales and could not be achieved by a single monospecies population. The degree of spatial structuring in the gut microbiota varies across length scales: at a macroscale of hundreds of micrometers, bacteria cluster into distinct habitats, whereas at a scale of micrometers intermixing of community members has been observed^3,10,11^.

The resilience of microbiomes, defined as the capacity to recover from perturbations, is strongly linked to microbial diversity. Indeed, a reduction in microbial diversity of the human gut microbiome is associated with multiple diseases, suggesting that a high-dimensional and functionally heterogeneous ecosystem promotes human health^12^. Understanding the molecular and ecological factors influencing the stability and resilience of the gut microbiota has implications for the development of targeted interventions to modulate system behaviors. Central to this problem is inferring unknown microbial interactions and developing tools to predict temporal changes in community behaviors in response to environmental stimuli.

The gut microbiota is composed of hundreds of bacterial species, the majority of which span the *Firmicutes*, *Bacteroidetes* and *Actinobacteria* phyla^13^. Constituent strains of the gut microbiota have been shown to persist over decades, demonstrating that community membership is stable as a function of time^14^. Perturbations to the system such as dietary shifts or antibiotic administration can shift the operating point of the gut microbiota to an alternative state^15^. While the identities of the organisms and microbial co-occurrence relationships across individuals have been elucidated^16^, we lack a quantitative understanding of how microbial interactions shape community assembly, stability and response to perturbations. For example, the molecular mechanisms that enable coexistence of the dominant phyla *Firmicutes* and *Bacteroidetes* as well as dynamic shifts in the *Firmicutes-to-Bacteroidetes* ratio, an indicator of human health^17,18^ are not well understood^8^.

The balance between cooperation and competition among constituent organisms dictates community-level properties such as metabolic activities and stability. Negative interactions have been shown to dominate microbial interrelationships in synthetic aquatic microcosms^19^. However, the prevalence of competition and cooperation in microbial communities occupying other diverse environments remains elusive. Direct negative interactions in microbial consortia can originate from competition for resources or space, biomolecular warfare or production of toxic waste products^20^. Positive interactions can stem from metabolite interchange or detoxification of the environment. The plasticity of pairwise microbial interactions can be augmented by conditional modulation by a third organism (higherorder interactions)^21^. Ecological driver species, which exhibit a large impact on community structure and function, represent key nodes in the network that could be manipulated for achieving maximal control of community behaviors^22^.

Deciphering microbial interactions is a challenging problem and key step towards understanding the organizational principles of microbial communities. Computational models at different resolutions can be used to infer microbial interaction networks^23^. Dynamic computational models can be used to predict and analyze temporal behaviors including response to perturbations and community assembly. Further, tools from dynamical systems theory can be used to analyze system properties including stability and parameter sensitivity^24^. A multitude of uncertainties in model structure and unknown parameters precludes the construction of mechanistic models that represents the bipartite network of metabolite-mediated interactions in microbial consortia. Generalized Lotka-Volterra (gLV) is a reduced complexity ordinary differential equation model that represents microbial communities with a limited number of parameters that can be deduced from properly collected time-series data.

Here we develop a systematic modeling and experimental pipeline to dissect the microbial interactions shaping the assembly of a synthetic community composed of 12 diverse species spanning major phyla of the human gut microbiota including *Bacteroidetes, Firmicutes, Actinobacteria*, and *Proteobacteria*. Time-resolved measurements of monospecies and pairwise assemblages were used to train a predictive dynamic computational model of multi-species communities. The network interaction pattern revealed influential and ecologically responsive organisms and was used to define the roles of constituent community members. *Bacteroidetes* exhibited an overall negative impact on the community, whereas specific members of *Actinobacteria* and *Firmicutes* displayed numerous cooperative inter-relationships. Pairwise topologies linked by positive and negative inter-relationships exhibited a broad parameter regime for stable coexistence in the model, providing insights into the networks that promote coexistence among phyla in the gut microbiota. The model showed that pairwise consortia that displayed history-dependent responses of initial species proportions were enriched for negative interactions and this behavior frequently did not arise from bistability, highlighting that community assembly can occur over a broad range of timescales. The metabolic capabilities of monospecies were elucidated using exo-metabolomics profiling and these data pinpointed a set of metabolites predicted to mediate positive and negative interactions. However, the metabolite profiles failed to forecast influential organisms modulating community assembly. Together, these results show that combinations of pairwise interactions can represent the composite behaviors of multi-species communities and these networks can realize a diverse repertoire of qualitatively distinct community responses.

## RESULTS

### Probing the temporal behaviors of monospecies and pairwise assemblages

We aimed to dissect the microbial interactions influencing community assembly in a reduced complexity model gut community. To this end, a synthetic ecology encompassing prevalent human-associated intestinal species *Bacteroides thetaiotamicron* (BT), *Bacteroides ovatus* (BO), *Bacteroides uniformis* (BU), *Bacteroides vulgatus* (BV), *Blautia hydrogenotrophica* (BH), *Collinsella aerofaciens* (CA), *Clostridium hiranonis* (CH), *Desulfovibrio piger* (DP), *Eggethella lenta* (EL), *Eubacterium rectale* (ER), *Faecalibacterium prausnitzii* (FP) and *Prevotella copri* (PC) was designed to mirror the functional and phylogenetic diversity of the natural system (**Fig.1a**) ^25^ These species have been shown to contribute significantly to human health and are implicated in multiple human diseases^26–31^ (**Supplementary Table I**).

**Figure 1.**
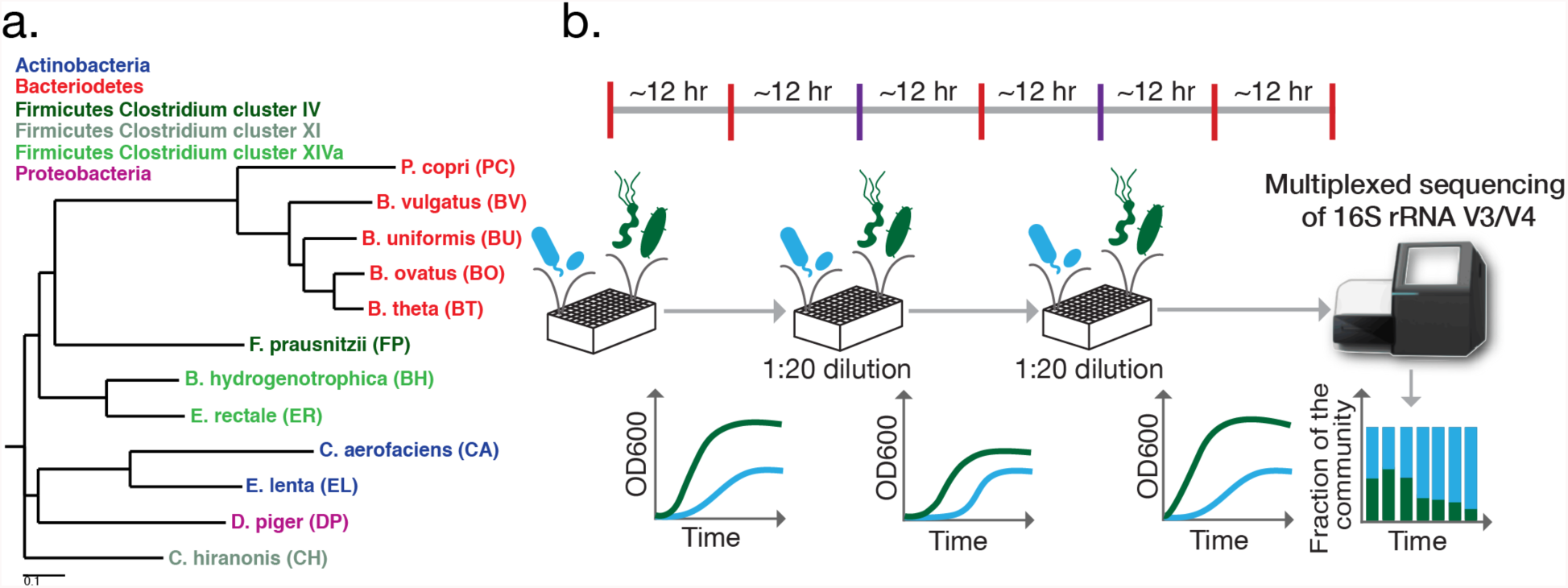
Experimental design for high-throughput characterization of synthetic human gut microbiome consortia. **(a)** Phylogenetic tree based on the 16S rRNA V3/V4 region of a 12-member synthetic ecology spanning the major phyla in the gut microbiome including *Actinobacteria*, *Bacteroidetes*, *Firmicutes* and *Proteobacteria*. **(b)** Schematic of the experimental design for this study. Species were combined at a 1:1 or 19:1 initial ratio based on absorbance measurements at 600 nm (0D600) into microtiter plates using liquid-handling robotic manipulation. Approximately every 12 hr, samples were collected for multiplexed 16S rRNA gene sequencing (red and purple bars, top). Relative abundance was measured using multiplexed 16S rRNA gene sequencing of the V3/V4 region using dual-indexed primers compatible with an Illumina platform (stacked bar plot, bottom right). To mirror periodic fluctuations in nutrient availability and monitor community assembly over many generations, serial transfers were performed in approximately 24 intervals (purple bars, top) by transferring the communities into fresh media using a 1:20 dilution. In parallel, time-resolved 0D600 measurements of monospecies and consortia were performed.

Synthetic assemblages were arrayed in microtiter plates in an anaerobic chamber using an automated liquid handling procedure (see Materials & Methods). The communities were serially transferred at 24 hr intervals to monitor community assembly over many cell generations and to reflect recurrent temporal perturbations to the gut microbiota such as diet and colonic transit time (**Fig. 1b**). Multiplexed 16S rRNA gene sequencing was performed in approximately 12 hr intervals to elucidate the temporal variations in community structure. The relative abundance for each species was computed as the sum of the read counts for each organism divided by the total number of reads per condition (see Materials & Methods). Since model construction is aided by absolute abundance information^32,33^, the total biomass of the communities was monitored approximately every 30 min using absorbance at 600 nm (OD600).

To infer microbial interactions, time-resolved measurements of monospecies and all pairwise communities (66 total) were performed using a 1:1 initial abundance ratio based on 0D600 values (PW1 dataset, **Fig. S1**). Monospecies growth and community composition were measured based on absorbance at 600 nm (OD600) and multiplexed 16S rRNA gene sequencing, respectively (see Materials & Methods). The monospecies displayed a broad range of growth rates, carrying capacities and lag phases (M dataset, **Fig. S2**). Pairwise consortia exhibited diverse growth responses and dynamic behaviors including coexistence or single-species dominance (**Fig. S1**). The distribution of absolute abundance of each species across communities in PW1 revealed changes in growth in the presence of a second organism (**Fig. S3**). Absolute species abundance was normalized to the monospecies carrying capacity to evaluate relative changes in the baseline fitness of each organism in the presence of second species. *Bacteroides* displayed unimodal distributions with a mean approximately equal to the monospecies carrying capacity, indicating the majority of species did not significantly modify *Bacteroides* fitness. Specific organisms such as PC, FP and DP exhibited bimodal or long-tail distributions, demonstrating that growth was significantly altered in the presence of specific organisms.

To further probe the dynamic responses of pairwise consortia, a set of 15 consortia (**Fig. 1b**) inoculated at two different initial species proportions (initial conditions) based on OD600 values (95% species A, 5% species B and the second wherein these percentages were reversed) were characterized using our experimental workflow (PW2 dataset, **Fig. S4**). The community behaviors in PW2 was classified into the following categories: (1) single species *dominance*; (2) *stable coexistence* wherein both species persisted above a threshold in the community for the duration of the experiment; (3) *history-dependence* whereby communities inoculated using distinct initial species proportions mapped to different community structures, or (4) *other* for communities that did not quantitatively satisfy the relative abundance thresholds for cases 1-3 (**Fig. S5**). A subset of the communities classified in the *other* category displayed weak history-dependent response potentially attributed to variability among biological replicates. The qualitative behaviors of the remaining 51 pairwise communities were classified based on community structure at an initial (t = 0) and final (t = 72 hr) time point using the PW2 experimental design wherein the organisms were inoculated using two distinct initial conditions (95% species A, 5% species B and the reciprocal percentages, **Fig. S5a**). Our results demonstrated that approximately 50%, 24% and 12% of pairwise communities displayed dominance, stable coexistence and history-dependence, respectively (**Fig. S5b**).

### Construction of a dynamic computational model of the community

A generalizable modeling framework was developed to infer parameters from time-series measurements of relative abundance and total biomass (OD600). The generalized Lotka-Volterra (gLV) model represents microbial growth, intra-species interactions and pairwise interspecies interactions and can be used to predict the dynamic behaviors of the community and analyze system properties such as stability and parameter sensitivity. To minimize overfitting of the data, a regularized parameter estimation method was implemented that penalized the magnitude of the parameter values (see Materials & Methods). Three training sets were evaluated based on predictive capability: (T1) M; (T2) M, PW1; (T3) M, PW1, PW2. A range of regularization coefficient values (*λ*) was scanned to balance the goodness of fit to the training sets and degree of sparsity of the model (**Fig. S6**). The parameterized gLV model trained on T3 captured the majority of pairwise community temporal responses (**Fig. 2a,b**). However, the model did not accurately represent the dynamic behaviors of a set of communities including BH,EL; PC,CA; BO,CH; ER,BH and PC,BH.

**Figure 2.**
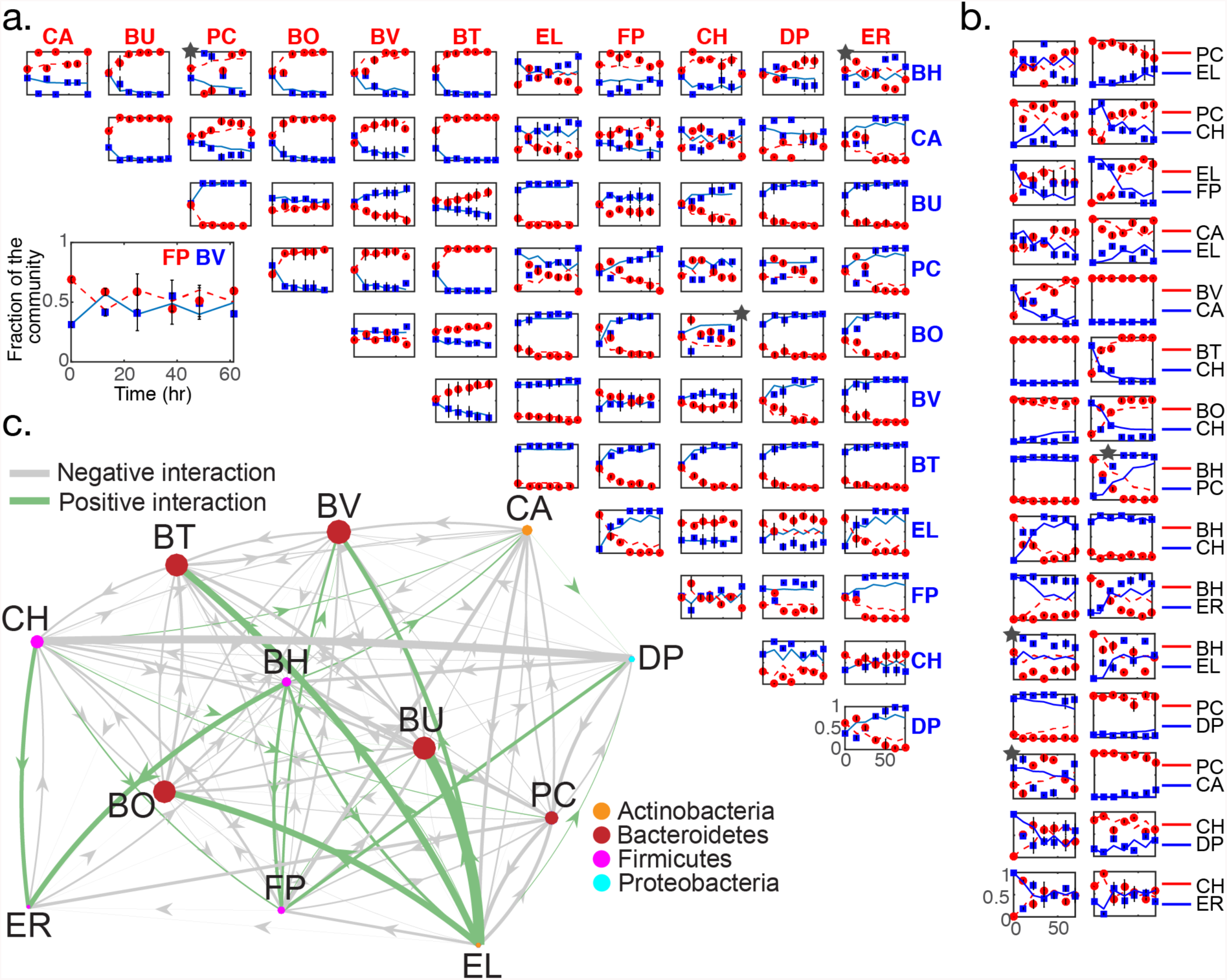
Model training of generalized Lotka-Volterra (gLV) to time-resolved measurements of monospecies and pairwise assemblages. **(a)** Relative abundance as a function of time for all pairwise communities. Experimental measurements and model fits are represented as data points and lines, respectively. The model was trained on T3 (M, PW1 and PW2). In each subplot, time and species relative abundance is displayed on the x-and y-axis, respectively. Stars denote datasets with a sum of mean squared errors greater than a threshold value of 0.15. Error bars represent 1 s.d. from the mean of at least three biological replicates. **(b)** Temporal changes in relative abundance of a selected set of pairwise assemblages inoculated at 5% species A, 95% species B or 95% species A, 5% species B based on OD600 values. Time and relative abundance are represented by the x and y-axis, respectively. Data points and lines represent experimental measurements and model fits, respectively. Error bars represent 1 s.d. from the mean of at least three biological replicates. **(c)** Inferred microbial interaction network based on model training on T3. Gray and green edges denote negative (α_ij_ < 0) and positive (α_ij_ > 0) interactions and the edge width represents the magnitude of the inter-species interaction coefficient. Node size is proportional to the monospecies steady-state abundance (x_e_ = -μ_i_α_ii_^-1^). To highlight significant interactions, inter-species interaction coefficients with a magnitude less than 1e-5 were not displayed.

Thresholding the magnitude of the inter-species interaction coefficients using a value of 1e-5 yielded a densely connected network whereby 77% of species pairs exhibited an interaction. Of these interactions, 61% and 16% were positive and negative, respectively (**Fig. 2c**). *Bacteroides* (BO, BV, BU and BT) displayed a net negative impact on the network, whereas EL, BH and CH positively stimulated a large number of species (**Fig. S7a**). Pairwise networks were enriched for amensalism (-/0, 36%), competition (-/-, 33%) and predation (+/-, 26%) (**Fig. S7b**). FP was the recipient of five positive interactions, suggesting that the fitness of FP is coupled to the composition of the community (**Fig. S3**). To determine the contribution of positive modulators to FP abundance, we examined a 6-member gLV model composed of FP, BH, BU, BV, CH and DP. The combined set of five positive inter-species interactions was required to alter FP abundance by >2-fold and single and dual inter-species interactions yielded a moderate increase in FP abundance at 72 hr (**Fig. S7c**). Therefore, FP represents an ecologically responsive organism that is significantly enhanced by the presence of multiple organisms in the community. Corroborating this notion, FP exhibited significant variability in absolute abundance across PW1 communities and frequent coexistence with other organisms (**Fig. S3, S5**). By contrast, BH (10 incoming interactions) and CH (9 incoming interactions) were negatively impacted by the largest number of species in the community. Hierarchical clustering of the gLV interaction coefficients demonstrated that phylogenetic relatedness was not a dominant predictor of microbial inter-relationships with the exception of *Bacteroides*, which exhibited a similar interaction pattern (**Fig. S8**). Therefore, evolutionary similarity was not a major variable structuring the microbial interaction network^34,35^.

### Model validation using informative multi-species assemblages

To validate the model, time-resolved measurements of relative abundance of the full (12-member) and all single-species dropout communities (11-member consortia, **Fig. 3a**) and total community biomass (**Fig. S9**) were performed. To determine if the synthetic community could recapitulate the abundance pattern observed across human gut microbiome samples, we compared the abundance profiles in the synthetic community at 72 hr and median relative abundance across human samples. The median abundance of 7 species in the metagenomics sequencing data of human fecal samples and mean abundance in the synthetic community were correlated (ρ = 0.79, P < 0.05)^36^, suggesting that microbial growth parameters and interspecies interactions are major variables influencing community structure in the human gut microbiota (**Fig. S10**).

**Figure 3.**
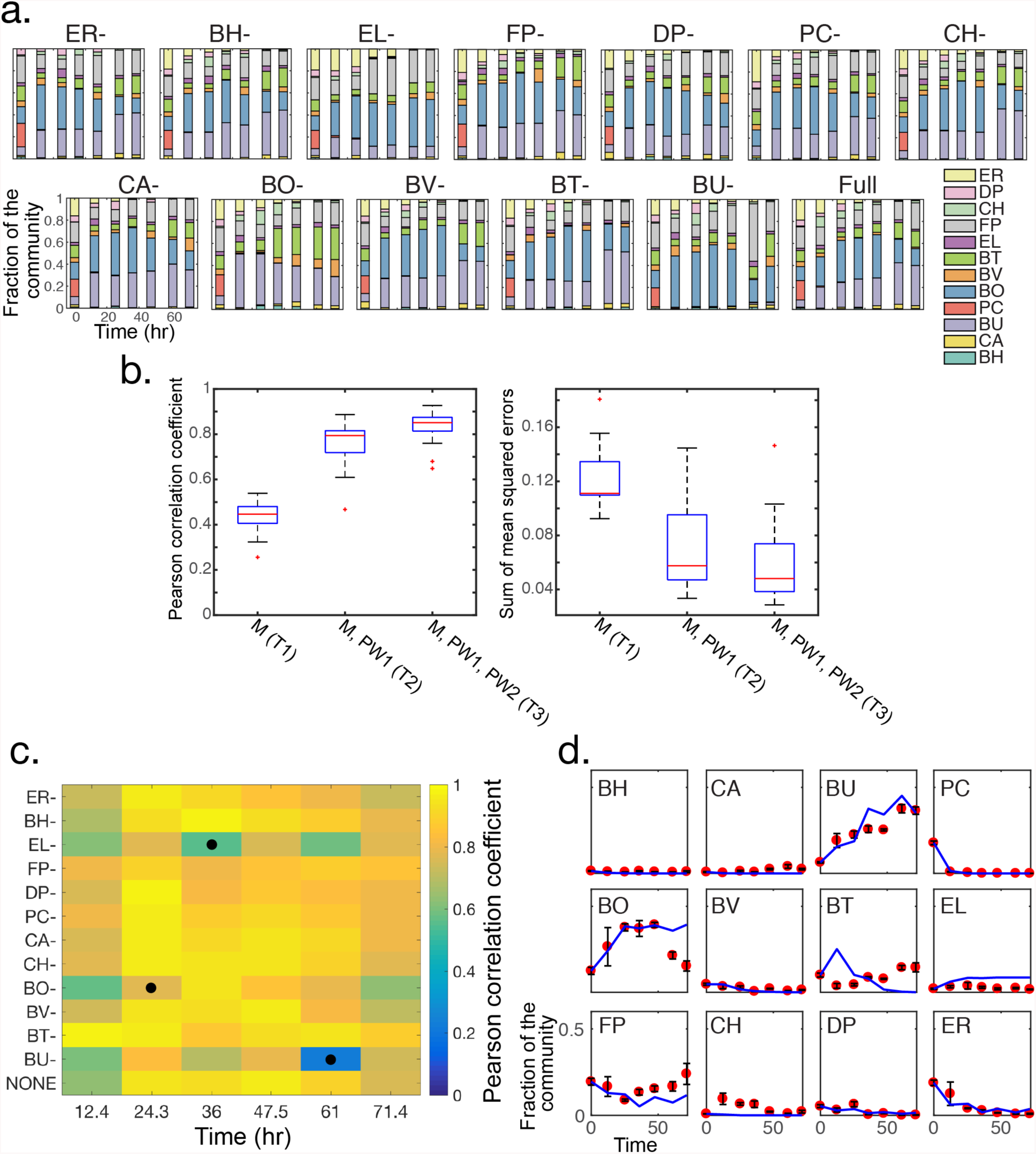
Validation of the parameterized Lotka-Volterra model to time-series data of informative multi-species communities. **(a)** Stacked bar plots of 11 and 12-member multi-species communities including all single species dropouts (X-represents the absent organism in the community) and the full community (full). Time and relative abundance are represented on the x and y-axis of each subplot, respectively. The absent organism is designated at the top of each subplot. Colors represent different organisms in the community. **(b)** Box plots of the metrics used to evaluate the predictive capability of models trained on different datasets. Pearson correlation coefficient for three model training sets (left) including T1: monospecies (M); T2: M and pairwise 1 (PW1); T3: M, PW1 and pairwise 2 (PW2). Sum of mean squared errors for the T1, T2 and T3 training sets (right). On each box, the red line represents the median, the edges of the box are the 25th and 75th percentiles, the whiskers extend to the most extreme data points and the outliers are plotted as red crosses. **(c)** Heat-map of Pearson correlation coefficients across time (x-axis) and multi-species communities (y-axis). Pearson correlations were statistically significant (P-value < 0.05) except for the EL-at 36 hr (P = 0.062), BU- at 61 hr (P = 0.47) and BO- at 24.3 hr (P = 0.051). Circles (black) denote conditions that were not statistically significant. **(d)** Comparison between model prediction and experimental data for each organism in the full community as a function of time. On each subplot, the x and y-axis represents time and relative abundance of each species, respectively. Data points (red) and lines (blue) denote experimental data and model predictions. Species names are displayed in the upper left corner of each subplot. Error bars represent 1 s.d. from the mean of six biological replicates.

Characterization of the temporal variation in community structure in the presence and absence of each organism elucidated the contributions of single species to community assembly. The absence of EL, BO or BU significantly altered community assembly compared to the full community, whereas elimination of all remaining organisms did not notably change community structure as a function of time (**Fig. S11a**). Further, the EL-consortium (community lacking EL) exhibited a lower species diversity scored by Shannon equitability index E_H_ (see Materials & Methods) for the duration of the experiment compared to the full community, indicating that EL promotes community diversity (**Fig. S11b**). In contrast to the EL-community, the BO- and BU-consortia exhibited a transient decrease in community diversity and delayed recovery to the full community E_H_ set point by 72 hr. The EL-community exhibited a significantly lower carrying capacity compared all other multi-species assemblages, indicating that EL plays an important role in shaping community structure, fitness and diversity (**Fig. S9**). The impact of EL on community dynamics was disproportionately related to its stable low abundance in the full community, in contrast to the influential and highly abundant organisms BO and BU (**Fig. 3a**). The species impact score (species relative abundance in the full community at 72 hr plus the sum of the outgoing gLV inter-species interaction coefficients) was correlated to the difference in community structure for single-species dropouts (ρ = 0.7, P < 0.05, **Fig. S12**), suggesting that the inferred ecological network and relative abundance pattern could explain the contributions of organisms to multi-species community assembly.

The predictive capabilities of the parameterized gLV models were evaluated using the time-series measurements of multi-species communities. Validation metrics included the Pearson correlation coefficient between the model prediction and relative abundance measurements and the sum of mean squared errors of relative abundance across all species in each community (see Materials & Methods). The model trained on T1 (M) exhibited the lowest median Pearson correlation coefficient (ρ) of 0.45 (not statistically significant) and largest error compared to the models trained on T2 (M, PW1) and T3 (M, PW1, PW2) (**Fig. 3b**), demonstrating that monospecies growth parameters alone are not predictive of multi-species community assembly. The addition of PW1 to model training significantly increased the predictive capability of the model (median ρ = 0.79), highlighting that pairwise interactions are major variables driving community dynamics. The inclusion of PW2 (T3 training set) further improved the correlation (median ρ = 0.85) and decreased the median error, indicating that the dynamic responses of communities inoculated using distinct initial species proportions were informative for inferring model parameters.

The predicted steady-state abundance of monospecies based on the model and the fraction of each species in the full community at 72 hr were not correlated (ρ = 0.46, P = 0.13), corroborating the notion that monospecies growth parameters alone failed to forecast temporal changes in multi-species structures (**Fig. S13**). Whereas FP exhibited low monospecies fitness (**Fig. S2**), this organism displayed the second highest abundance level in the full community, which was consistent with a large number of positive incoming interactions present in the inferred gLV network (**Figs. 2c, S7a,c, S13**). By contrast, BV persisted at low abundance in the full community and exhibited high steady-state monospecies abundance, consistent with a large number of inhibitory incoming inter-species interactions (**Fig. S7a**). Therefore, the pattern of ecological interactions provided insight into significant deviations in species fitness in the absence and presence of the community.

To evaluate the predictive capability of the model, the quantitative relationship between the model and data was examined for each species as a function of time using the parameter estimate based on training set T3 (**Fig. 3c**). Across the majority of time points and multi-species consortia, the Pearson correlation coefficient between the model prediction and experimental data explained a high fraction of the variance and was statistically significant. While the model captured the temporal responses of the majority of species, the model did not reproduce the behavior of BT in the multi-species consortia and deviated at specific time intervals for BO and CH (**Fig. 3d**).

Incorporating additional data sets into the model training procedure could reduce the contributions of noise to parameter estimates and provide additional parameter constraints. To this end, the model was trained on M, PW1, PW2 and the time-resolved measurements of the full community (training set T4) and was validated on the set of single-species dropout communities (**Fig. S14**). The model predictive capability trained on T4 was improved compared to T3 and explained 81% of the variance (ρ = 0.9) on average of the multi-species community temporal responses. Model parameter sets based on T3 and T4 were highly correlated with the exception of a small number of microbial interactions (**Fig. S14b**). Approximately half (55%) of the interaction coefficients that were present or absent based on model training on T4 compared to T3 involved the ecologically influential species EL and BO (**Figs S14c, S11a**). In sum, the pairwise gLV model could accurately predict multi-species community dynamics and thus conditional interactions played a minor role in driving community assembly.

### History dependence and robust coexistence in pairwise assemblages

We analyzed the model to elucidate the origins of history-dependent behaviors in pairwise communities (**Fig. S5**). The inferred network based on the model revealed that 6 of the 8 pairwise communities that exhibited history-dependent behaviors were linked via mutual inhibition and the remaining two networks displayed unidirectional negative interactions (**Fig. 4a**). To probe the quantitative behaviors of the mutual inhibition network topology, we analyzed the parameter regime that generates history-dependence and bistability. Bistability is a property of a dynamical system whereby the system has two stable steady-states and can exhibit longterm dependence of the state of a system on its history, referred to as hysteresis. Bistability is a possible outcome of a gLV model of pairwise mutual inhibition^37^.

**Figure 4.**
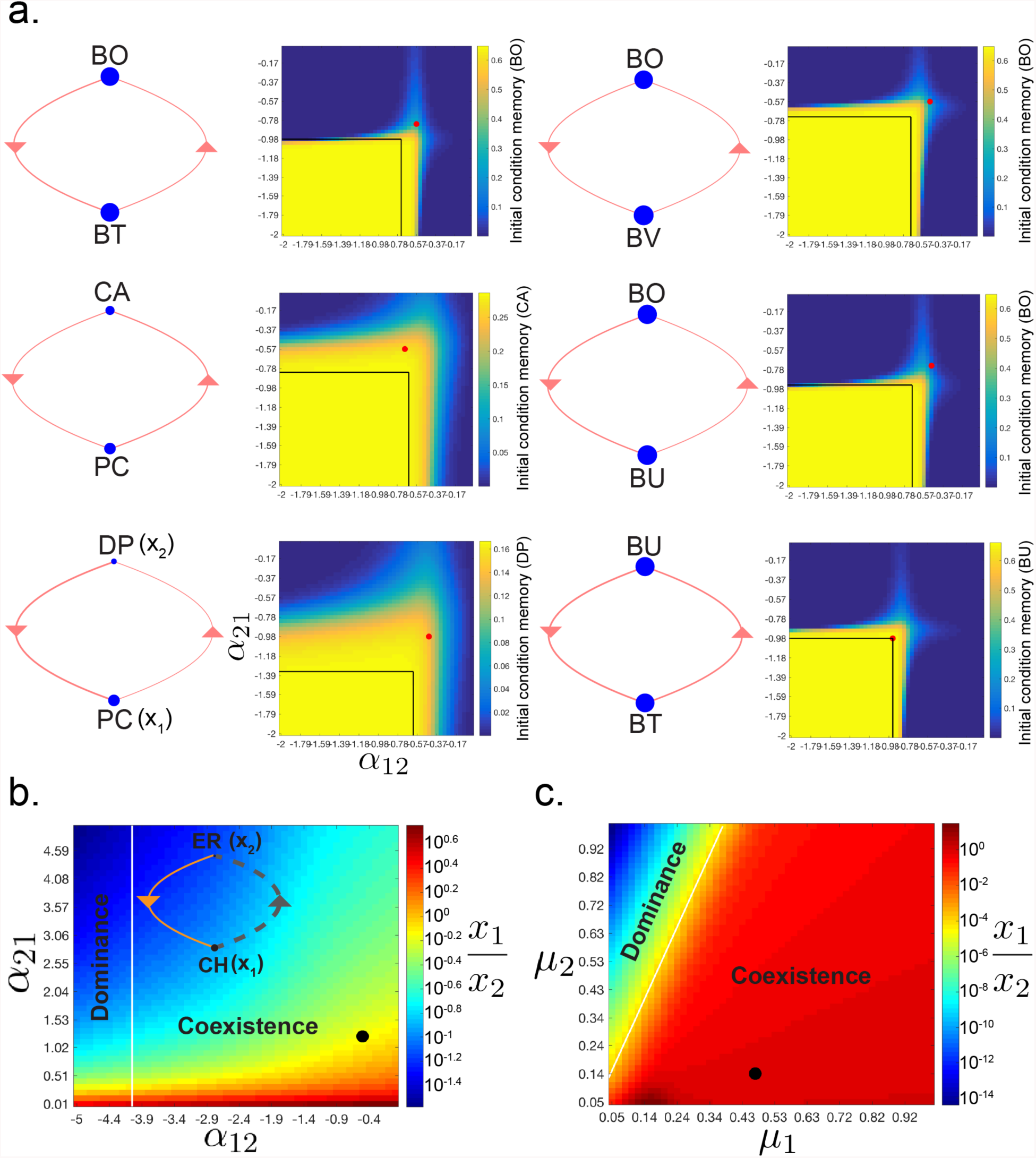
History-dependence and robust species coexistence in pairwise consortia motifs. **(a)**History-dependence can stem from negative inter-species interactions in pairwise communities. Six pairwise communities that experimentally displayed history-dependent behaviors (**Fig. S5b**) and are coupled by mutual inhibitory interactions in the inferred network. Network topology (left) and heat-map of history-dependent responses (right) across a range of inter-species interaction coefficient values. The line width and node size of the network diagram represents the magnitude of the inter-species interactions and steady-state monospecies abundance, respectively. Memory of initial conditions in the model is defined as the absolute value of the difference in species absolute abundance at 72 hr for communities initialized at two different conditions: x_1_ = 0.0158, x_2_ = 0.0008 or x_1_ = 0.0008, x_2_ = 0.0158 using a serial dilution experimental design (**Fig. 1b**). The black box denotes the set of α_12_ and α_21_ values yielding bistability in the model. The circle (red) indicates the inferred parameters based on training set T3. **(b)** Coupled positive and negative interactions can yield a broad parameter regime for species coexistence. Network diagram (inset) represents the magnitude, sign and direction of the inferred inter-species interactions between CH (x_1_) and ER (x_2_). Dashed (gray) and solid (orange) lines indicate a positive and negative interaction, respectively. The line width and node size denotes the magnitude of the interspecies interaction coefficients and steady-state abundance of the monospecies, respectively. Heat-map of the ratio of x_1_ (CH) to x_2_ (ER) at 72 hr as a function of the inter-species interaction coefficients α_12_ and α_21_. Initial conditions for simulations were x_1o_ = 0.0008, x_2o_ = 0.0158. The circle (black) indicates the inferred parameters for the CH, ER consortium based on training set T3. **(c)** Heat-map (right) of the ratio of x_1_ to x_2_ at 72 hr across a broad range of growth rate parameter values (μ_1_ and μ_2_). Initial conditions for simulations were x_1o_ = 0.0008, x_2o_ = 0.0158. The line (white) outlines the parameter regimes for coexistence and single-species dominance at steady-state. The circle (black) represents the inferred parameter values for the CH, ER consortium.

History-dependent dynamic behavior in the model was defined as the difference in species abundance following 72 hr for communities inoculated using two distinct initial species proportions (95% species A, 5% species B and the reciprocal condition) and does not require bistability. In 5 of the 6 mutual inhibition networks, the pairwise consortia were not operating within a bistable parameter regime, indicating that the observed history-dependent responses stemmed from a prolonged duration of time for community assembly. A range of inter-species interaction coefficient values beyond the bistable parameter regime displayed memory of initial conditions that persisted for at least 72 hr, indicating that mutual inhibition can generate protracted kinetic dependence of the states of a system on its history. The inferred parameters for the bistable BU, BT consortium were located on the boundary between monostability and bistability, suggesting that the system’s bistable behavior was not robust to parameter variations (**Fig. 4a**).

To investigate the physiological significance of such history-dependent behaviors, we examined the capacity for history-dependence across a broad range of simulated pairwise consortia serial dilution rates (**Fig. S15**). The rate of serial dilution represents colonic transit time, a major variable shown to modulate human gut microbiome composition, diversity and functions^38,39^. Pairwise communities BO,BT; DP,PC and CA,PC exhibited history-dependent responses up to 168 hr across a range of dilution rates due to strong inhibitory microbial interactions. Together, our modeling results demonstrate that the frequency of environmental shifts can promote history-dependent behaviors over physiological timescales (days) in monostable pairwise communities coupled by mutual or unidirectional negative interactions. Perturbations that shift the system’s operating point away from steady-state could augment the duration of time that the system displays history-dependence due to slow timescales for returning to a steady-state. Together, these results highlight that community assembly can evolve over long timescales, thus challenging a steady-state operating point assumption for the human gut microbiome^40^.

A subset of pairwise consortia exhibited stable coexistence wherein communities inoculated at distinct initial species proportions converged to a non-zero abundance level above a defined threshold that persisted for the duration of the experiment (**Fig. 2b, S5**). Stable coexistence whereby both species have non-zero abundance at steady-state is a possible outcome of the generalized Lotka-Volterra model^37^. To evaluate the robustness of coexistence across different network topologies, we analyzed a pairwise consortium composed of CH and ER that converged to an approximately equal abundance ratio as a function of time from two disparate initial conditions (**Fig. 2b**). The inferred pairwise network consisted of a positive and negative coupling between CH and ER (**Fig. 4b**). The positive and negative interaction topology yields a broader parameter regime of coexistence compared to mutual inhibition (**Fig. 4b,c, S16a,b**). The combination of positive and negative interactions establishes a negative feedback loop on the abundance of an organism that has higher monospecies fitness (e.g. CH), thus leading to stable coexistence with an organism that has lower monospecies fitness (e.g. ER) as a function of time.

Coexisting species pairs were linked by positive and negative interactions and mutual inhibition in 50% and 25% of cases, mirroring the disparity in parameter space that realized coexistence between the two distinct topologies (**Fig. 4b,c, S16**). The remaining species pairs that displayed coexistence encompassed unidirectional positive (12.5%), unidirectional negative (6.25%) and mutualism (6.25%). Several dominant phyla were connected by positive and negative interactions in the phylum-level interaction network, representing the average interspecies interaction coefficient for all organisms associated with a given phylum, suggesting that this topology promotes stable coexistence in the gut microbiota (**Fig. S17**).

### Analysis of model parameter constraint

Previous parameter estimation methods have failed to examine the uncertainty in parameter estimates^41,42^. Methods from Bayesian statistics can illuminate the uncertainty in the parameter values using the Posterior distribution, which represents the probability of the parameters given the data. To this end, the Metropolis Hastings Markov Chain Monte Carlo (MCMC) method was implemented to randomly sample from the posterior distribution (see Materials & Methods). The coefficient of variation (CV) of 99% of parameters was less than 0.5 for 200000 iterations, indicating that the parameters were constrained by the data (**Fig. S18a**). Parameters that were present or absent based on training on T4 compared to T3 exhibited a higher median CV value of 0.22 compared to the unchanged set that had a median CV equal to 0.16 (**Figs. S14b,c, S18b**). These data suggest that model training on T4 provided additional constraint for specific parameters that displayed larger uncertainty in parameter estimates.

The correlation between all parameter pairs in the model (12090 total combinations) was evaluated to determine parameter identifiability (**Fig. S18c**). Parameters that do not influence observable variables are non-identifiable due to practical or structural reasons and can be correlated with other parameters^43^ (**Fig. S18d,e**). Approximately 11% of parameter pairs displayed a Pearson correlation coefficient greater or less than 0.6 and -0.6, indicating that the majority of parameters could be distinguished. Correlated interaction coefficients greater or less than 0.6 and -0.6 were approximately 7-fold lower magnitude on average than parameters that exhibited higher identifiability (**Fig. S18f**). In the parameter estimation procedure, regularization will lead to the reduction of the magnitude of parameters that do not link to the observable outputs.

### Interrogating microbial environmental impact using conditioned media

The net environmental impact of a single species at a defined time point can be represented by conditioned media, which contains secreted metabolites and has been depleted for specific resources. Positive interactions may be indicative of transformations of media components into substrates that can be utilized by the recipient species or detoxification of the environment. Negative interactions may derive from depletion of key nutrients or production of toxic compounds. In some cases, multiple mechanisms can combine to yield a net positive or negative effect on growth. Environmental pH is a major variable that can influence microbial growth responses. In co-culture, the environmental pH may not change as significantly compared to monoculture growth due to differences in metabolite secretion and degradation in a community. For example, cross-feeding of metabolic by-products such as acetate in the gut microbiota is a prevalent mechanism that could alter environmental pH^44–46^.

We investigated whether the difference in the recipient species growth responses in the presence and absence of conditioned media from a source organism could be used to map microbial inter-relationships (**Fig. S19**). To this end, a conditioned media impact score R_CM_ was defined as the ratio of the cumulative sum of the recipient organism growth response for 30 hr in 75% conditioned media to unconditioned media. An R_CM_ > 1 or R_CM_ < 1 indicated a positive or negative influence of the source organism on the recipient organism. To evaluate the contribution of pH to the conditioned media growth responses, pH adjusted conditioned media was prepared by modifying the pH to match the value of the unconditioned media.

Several factors could lead to disagreements between R_CM_ and the gLV inter-species interaction coefficients, including, for example, a difference in metabolite utilization and secretion patterns of an organism in the presence of a second species^47^. Nevertheless, 75% of conditions were in qualitative agreement with the inferred gLV interaction coefficients (**Fig. S19**). Of the interactions that showed qualitative disagreement between conditioned media and inferred gLV interaction coefficients, 45% displayed low identifiability based on Metropolis-Hastings MCMC using Pearson correlation coefficient thresholds of -0.6 and 0.6, suggesting that model parameter uncertainties could also contribute to the observed inconsistencies. Together, these data corroborated the parameter estimation pipeline and showed that a high fraction of microbial inter-relationships could be qualitatively deciphered based on conditioned media responses.

### Elucidating metabolic capabilities of monospecies via exometabolomics

Metabolite interchange is a dominant mode of microbial interactions. To elucidate the bipartite structure of the metabolite and species network for the synthetic community, exo-metabolomics profiling of 97 major metabolites was performed on monospecies. The metabolite profiles were analyzed at an initial and final time point that occurred prior to 24 hr to mirror the community experimental design with the exception of DP due to insufficient accumulation of biomass for metabolomics measurements within the 24-hour period (**Fig. 1b, S20**). Relative changes in metabolite abundances were computed using the log2 fold change of the final and initial time point and a significance threshold of at least two-fold was applied to the data (**Fig. 5a,b**). We performed clustering analysis of the metabolite utilization and secretion bipartite networks to identify similarities in metabolite profiles. The clustering pattern did not the recapitulate the phylogenetic relationships, demonstrating that distantly related species can occupy similar resource utilization niches (e.g. BH and EL or FP and ER) and closely related species can utilize distinct resources (e.g. BH and ER) (**Fig. 1a**).

**Figure 5.**
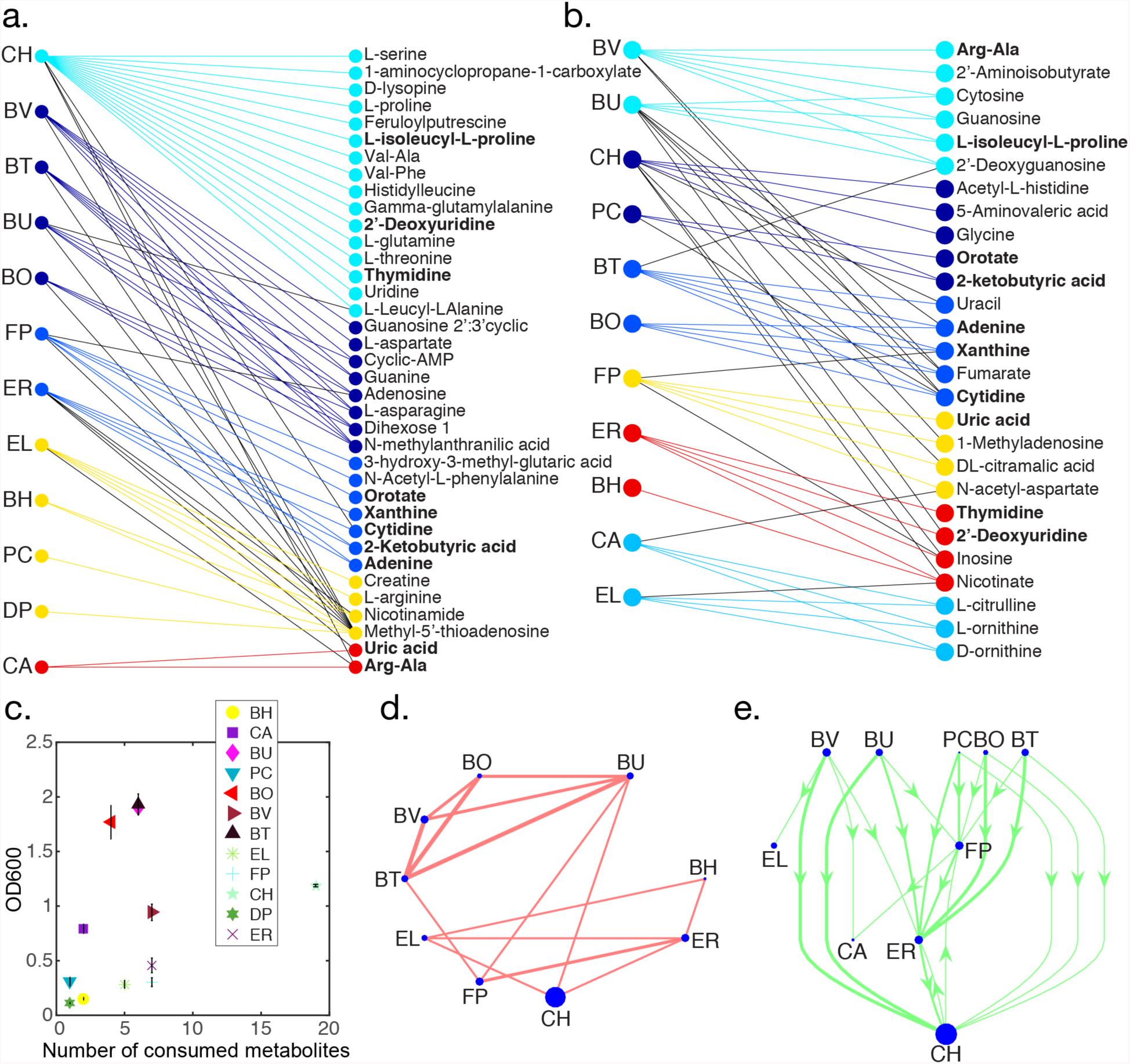
Exo-metabolomic profiling of major metabolites elucidated the metabolic capabilities of monospecies. **(a)** Bipartite network of species (left) and metabolites (right) for metabolites that decreased by at least two-fold compared to the abundance of each metabolite at the beginning of the experiment. Colors represent modules containing many overlapping interactions in the network. Network partitioning into modules was performed using BiMat^48^. Metabolites in bold were depleted and secreted by distinct organisms. **(b)** Bipartite network of species (left) and metabolites (right) for metabolites that increased in abundance by at least two-fold compared to the beginning of the experiment. Metabolites highlighted in bold were depleted or secreted by different species. **(c)** Scatter plot of the number of consumed metabolites using a threshold of two-fold vs. the OD600 value of the monospecies culture at the corresponding time point. Error bars represent 1 s.d. from the mean of three biological replicates. **(d)** Predicted resource utilization interaction network. Each edge represents at least two co-consumed metabolites and the edge width is proportional to the number of co-consumed metabolites. Node size is proportional to the total number of consumed metabolites for each species. **(e)** Predicted metabolite interchange network representing metabolites that were secreted or utilized by distinct organisms. Arrows point from the source species to the consumer organism. Node size and line width are proportional to the total number of secreted metabolites and number of predicted metabolite interactions, respectively. Species at the top and bottom of the network are primarily producers or consumers, respectively.

Our results showed a lack of correlation between the number of consumed metabolites and the total biomass produced by each monospecies at the corresponding time point. CH consumed the largest total number of metabolites in comparison to other organisms, thus representing a hub in the metabolite utilization network (**Fig. 5c**). The total biomass produced by CH was not proportional to the total number of consumed metabolites or sum of log2 fold changes in utilized metabolites, suggesting that CH could be funneling energy towards cellular processes beyond biomass (**Figs. 5c, S21a**). Corroborating these results, CH exhibited a large number of positive and negative outgoing edges in the inferred inter-species interaction network (**Fig. S7a**).

The sum of log2 fold changes in metabolite secretion was correlated with total monospecies biomass (R_s_ = 0.83, P < 0.002), relative abundance in the full community at 72 hr (R_s_ = 0.66, P < 0.05) and the number of outgoing negative interactions for each organism (R_s_ = 0.65, P < 0.05) based on inferred ecological network, where R_s_ represents the Spearman rank correlation coefficient (**Fig. S21b,c,d**). These data suggest that metabolite secretion was a better predictor of species fitness and negative ecological interactions compared to the metabolite utilization pattern. The resource co-utilization network pinpointed significant resource competition among *Bacteroides* for a set of core metabolites (**Fig. 5d**). A network of predicted metabolite interchange illuminated producers (*Bacteroidetes*), consumers (EL and CA) and species that played dual roles (FP, ER and CH). Metabolites predicted to mediate the largest number of pairwise negative interactions via resource competition included methyl-5’-thioadenosine (55 pairs), N-methylanthraniic acid (6 pairs) and dihexose (3 pairs) (**Fig. S22a**). Metabolites implicated in three or more positive interactions due to predicted metabolite cross-feeding encompassed cytidine, adenine and 2-ketobutyric acid (**Fig. S22b**). FP was predicted to utilize several metabolites produced by BU, BO, PC, BT and CH, suggesting a potential molecular basis of positive modulation by BU and CH in the inferred inter-species interaction network (**Fig. 2c, Fig. S7a,c**). However, the metabolite profiles failed to predict the influential role of EL in mediating community assembly, fitness and diversity (**Fig. 5d,e**).

## DISCUSSION

Developing the capabilities to predict microbial community dynamics in response to environmental stimuli is a first step towards elucidating the organizational principles of microbial communities and devising strategies for precisely manipulating ecological properties. The discovery of significant microbial inter-relationships and ecological driver species in the network can be exploited as novel control parameters for microbiomes. To this end, we developed a generalized parameter estimation pipeline to build predictive dynamic models of microbial communities complementary to previously published methods^49^. In contrast to statistical network models, dynamic frameworks can be used to extract mathematical principles and probe system properties such as ecological stability, history-dependence and response to perturbations. Further, the inferred network can be used to define ecological roles for each species and model could be harnessed as a predictive tool for designing sub-communities with desired properties. We capitalized on methods from Bayesian statistics to go beyond a single parameter estimate to evaluate the uncertainty in parameters given the data. Future work will harness this information for experimental design by iteratively guiding the selection of informative experiments to reduce parameter uncertainties and thus enhance the predictive capabilities of the model.

Our results substantiate the notion that pairwise interactions dominate multi-species ecological dynamics^50^. Previous work showed that pairwise phenomenological models of low dimensional assemblages (2-3 species) trained on an interval of time of Monod-based community models failed in some cases to predict future dynamic behaviors^51^. The mechanistic models considered in this study involved a limited number of metabolites, whereas microbes likely interact via a high-dimensional vector of metabolites. Future work will explore the capability of pairwise models to recapitulate the dynamics of mechanistic models that capture such complexities. Here we show that pairwise interactions can realize diverse behaviors encompassing history-dependence, coexistence and single-species dominance. Combinations of such interactions in multi-species assemblages can yield a diverse repertoire of dynamic behaviors and realize systems-level properties including stability, resistance to invasion and resilience to disturbances^52–54^.

We find that negative interactions can promote history-dependent responses over physiological timescales in response to environmental perturbations, demonstrating that the timescales of community assembly are contingent on network couplings and growth parameters. As such, dynamic frameworks are essential for dissecting and forecasting system behaviors away from steady-state. The steady-state assumption for gut microbiome composition is not valid if the timescales of environmental perturbations that steer the system away from steady-state occur faster than the time required to converge to a steady-state^40^. Indeed, inputs to the gut microbiota such as dietary shifts may steer the system away from a steady-state^55^, thus forcing the system to operate in a regime outside of steady-state due to prolonged timescales of community assembly.

We interrogated a synthetic ecology composed of prevalent human-associated intestinal species that play major roles in human health and disease (**Supplementary Table I**). Our systematic approach to interrogate community assembly rigorously defined the ecological contributions of community members based on the inter-species interaction patterns. For example, the pairwise network revealed hubs for negative (*Bacteroidetes*) and positive interactions (EL, CH and BH). A top-down community assembly approach to investigate single-species dropouts pinpointed influential organisms that shape multi-species community assembly including EL, BO and BU. A positive and negative interaction topology that can realize robust species coexistence linked representative species from the major phyla *Bacteriodetes*, *Firmicutes* and *Actinobacteria*. *Bacteroidetes* and *Firmicutes* have unique and complementary metabolic specializations in the gut microbiota^8^, consistent with the numerous positive interactions deciphered between members of these phyla. *Bacteroides* and *Prevotella* have been shown to be anticorrelated across individuals^56,57^. A recent study demonstrated that gnotobiotic mice colonized with BT and PC exhibited lower absolute abundance of both species compared to mono-colonized gnotobiotic mice, suggesting an inhibitory interaction^58^. In line with these results, BT excluded PC in pairwise experiments (**Fig. 2a, S5a**) and the inferred network illuminated a negative outgoing interaction from BT to PC (**Fig. 2c**). A previous study showed that BT negatively influences BV in metagenomics time-series data from one individual, corroborating a negative outgoing interaction from BT to BV in the inferred gLV network (**Fig. 2c**)^59^. Future work will illuminate the molecular mechanisms that promote stable coexistence and dynamic shifts in the proportions of key organisms in the gut microbiota in response to environmental perturbations.

Exo-metabolomics profiling identified CH as a hub for metabolite consumption and BU and CH as major producers of metabolites in the community. Indeed, CH had the potential for significant environmental impact via utilization and secretion of a broad repertoire of metabolites. However, CH was low abundance in the full community and a community lacking CH did not exhibit significant changes in community dynamics, fitness or diversity compared to the full community, suggesting that a large number of inhibitory incoming interactions precluded CH from playing an influential role in community dynamics (**Figs. 3d, S7a, S9, S11a**). Therefore, ecological inter-relationships can limit or enhance an organism’s potential for environmental impact in the context of a community.

The patterns of central metabolite consumption and secretion failed to predict influential species in community assembly, indicating a complex mapping between metabolite interchange and microbial interactions. For example, EL--a major driver species in the community--performed transformations on a moderate number of metabolites that were not implicated in metabolite interchange, whereas the metabolite hub CH did not significantly shape community assembly. Beyond central metabolites, secondary metabolites or signaling molecules could contribute to the observed ecological relationships. Metabolite secretion as opposed to utilization was correlated to the number of negative inter-species interactions, suggesting that negative interactions may derive from mechanisms beyond resource competition such as biomolecular warfare or production of toxic metabolic by-products. Negative interactions could lead to funneling of intracellular resources towards non-metabolic cellular processes such as stress, thus contributing to the lack of correlation between ecological interactions and resource utilization.

Microbial interactions have been probed in several synthetic ecologies of varying complexity including 18 *Streptomyces* strains^60^, 4-member freshwater isolates^61^ and 8 soil bacterial isolates composed of 6 strains from the *Pseudomonas* genus^50^. A significant fraction of the *Streptomyces* pairwise communities displayed frequent history-dependent and rare coexistence behaviors, which could be attributed to numerous mutual inhibitory pairwise interactions^60^. By contrast, history-dependent responses were not detected in the synthetic consortium of 8 soil isolates and coexistence was the prevalent community behavior^50^. On timescales of minutes, members of the freshwater isolate community did not display negative interactions that diminished cellular redox activity^61^. Here, we interrogated the pairwise community dynamic behaviors in a 12-member anaerobic community spanning four distinct phyla. This community showed diverse pairwise behaviors including single-species dominance, coexistence and history-dependence. Lower energy yields from anaerobic metabolism, which requires the concerted activities of distinct community members to achieve chemical transformations, may lead to differences in the pattern of ecological inter-relationships in anaerobic vs. aerobic communities. Such variations in the molecular mechanisms driving microbial interactions across distinct environments ranging from soil to the human intestinal system can manifest as differences in community-level properties including diversity, stability and dynamic responses to perturbations.

Positive interactions were observed frequently in the synthetic human gut microbiome community^19^. Indeed, the influential species EL positively impacted many species in the network, demonstrating that cooperation, as opposed to competition, can significantly influence community assembly, fitness and diversity. Therefore, our results underscore the role of cooperation in promoting ecological diversity in contrast to a previous study that showed negative interactions augment community diversity and stability in models of ecological networks^54^. Future work will elucidate how the plasticity of pairwise interactions in response to changeable environments mediates community assembly and stability in higher-dimensional microbiomes that mirrors the complexity of the natural system.

## MATERIALS AND METHODS

### Starter culture inoculations

Cells were cultured in an anaerobic chamber (Coy Lab Products) using mixed gas tanks containing 85% N_2_, 5% H_2_ and 10% CO_2_. Starter cultures for community measurements were inoculated from 200uL single-use 25% glycerol stocks into 15mL of Anaerobic Basal Broth (ABB) media (Oxnoid) in an anaerobic chamber and incubated at 37°C without shaking. To compare strains in similar growth phases that displayed variability in the duration of lag phases, the strains were partitioned into slow and fast growth categories inoculated 41 hr or 16 hr prior to the beginning of the experiment, respectively. Strains in the slow growth category included *B. hydrogenotrophica* (BH), *F. prausnitzii* (FP), *C. aerofaciens* (CA), *P. copri* (PC), *E. rectale* (ER), and *D. piger* (DP). Fast growing strains encompassed *B. uniformis* (BU), *B. vulgatus* (BV), *B. thetaiotaomicron* (BT), *B. ovatus* (BO), *C. hiranonis* (CH), and *E. lenta* (EL).

### Microbial community culturing

Each constituent strain was inoculated at 0.01 OD600 in microbial communities unless otherwise noted. In multi-species assemblages, the total initial OD600 was 0.01 OD600 x *n*, where *n* represents the number of strains in the community. For PW1, each component was normalized to OD600 of 0.01 and then mixed in equal proportion. For PW2, the major and minor strains were inoculated at 0.0158 and 0.0008 OD600, respectively. The microbial communities were arrayed using a liquid-handling robot (Biomek) into 96-well sequencing plates covered with a gas-permeable seal (Breath-Easy) and incubated at 37°C without shaking. In parallel, aliquots of the communities were transferred into a 384-well absorbance plate and grown in a F200 Infinite plate reader (Tecan) for OD600 measurements at 30 min intervals. Samples from the sequencing plate were collected approximately every 12 hr for a total of 72 hr. At each time point, samples were mixed and 400 μL was transferred to a 96-well collection plate. The collection plate was centrifuged at 4000 RCF for 10 min and 380 μL of the supernatant was removed with a multichannel pipette. Serial transfers were performed at 24 hr intervals into fresh media using a 1:20 dilution. Species diversity was scored by the Shannon equitability index E_H_ where *E_H_* = *H* ln *S*^−1^ and 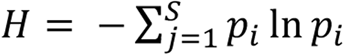. Here, S, H and p_i_ denotes the number of species in the community, Shannon diversity index and relative abundance of the *i*th species, respectively.

### Genome extractions

Genomic DNA (gDNA) extractions were performed using the QIAamp 96 DNA QIAcube HT Kit (Qiagen) with minor modifications including an enzymatic lysis pre-treatment step and the use of a vacuum manifold to perform column purification steps. Enzymatic lysis was performed as follows: cell pellets were re-suspended in 180 μL of enzymatic lysis solution containing 20 mg/mL lysozyme (Sigma-Aldrich), 20 mM Tris-HCl pH 8 (Invitrogen), 2 mM EDTA, and 1.2% Triton-X-100. Samples were incubated at 37°C for 30 min with shaking. Following the initial incubation, 4 μL of 100 ng/μL RNAse A (Qiagen) was added and samples were incubated at room temperature for ~1 min prior to administering 125 μL of proteinase K to buffer VXL (Qiagen). Samples were incubated for an additional 30 min at 56°C with shaking, 325 μL of ACB buffer was added and the samples were transferred to a 96-well column plate for purification. Samples were washed with 600 μL of AW1, AW2, and ethanol (Sigma-Aldrich). Following the ethanol wash, samples were allowed to dry for approximately 5 min. Finally, the gDNA samples were eluted using AE buffer pre-warmed to 56°C into a 96-tube rack plate and stored at -20°C.

### Illumina primer design, library preparation and sequencing

Dual-indexed primers were designed for multiplexed next-generation amplicon sequencing on Illumina platforms. Each 90-99 base pair (bp) forward and reverse primers consisted of an indexed 5’ Illumina adaptor, heterogeneity spacer^62^ and 3’ annealing region to amplify 466 bp of the V3-V4 variable region of the 16S rRNA gene. The set of 64 unique forward (6 bp) and reverse (8 bp) indices allowed multiplexing of 1536 samples per sequencing run. Oligonucleotides (Integrated DNA Technologies) were arrayed into 96-well plates using a stock concentration of 1 μM.

Following gDNA extraction, gDNA concentrations were quantified using the Quant-iT dsDNA High Sensitivity kit (ThermoFisher) and normalized to approximately 3 ng/μL. PCR amplification of the V3/V4 region of the 16S rRNA gene was performed with Phusion High Fidelity DNA-polymerase (NEB) for 18-25 cycles using 0.05 μM of each primer. PCR amplicons were pooled by plate (96 conditions), purified (Zymo Research) and quantified using the Quant-iT dsDNA High Sensitivity kit. The samples were normalized to the lowest sample concentration and then combined in equal proportions to generate the library. The library was quantified prior to loading using quantitative real-time PCR (KAPA Biosystems) on a CFX96 real time PCR detection system (Bio-Rad). Following amplification, the library was diluted to 4.5 nM and loaded on the Illumina MiSeq platform for 300 bp paired-end sequencing.

### Data analysis pipeline for 16S rRNA gene sequencing

A reference database containing the V3/V4 16S rRNA gene sequences was constructed by assembling consensus sequences based on next-generation sequencing of monospecies cultures. To process the sequencing data, the MiSeq Reporter software demultiplexed the indices and generated the FASTQ files using the bcl2fastq algorithm. Custom Python 2.6.6. scripts were used for subsequent data processing steps and are available for download at Github (https://github.com/ryanusahk/NextGenSequencingScriptsRH). First, paired-end reads are merged using PEAR (Paired-End reAd mergeR) v0.9.0^63^. The global alignment tool in USEARCH v8.0 mapped each sequence to the reference database. A 97.5% alignment threshold was implemented to distinguish the closely related species *B. thetaiotaomicron* and *B. ovatus.* Relative abundance was computed by summing the read counts mapping to each organism divided by the total number of reads per condition. The data was exported for analysis in MATLAB (Mathworks).

### Conditioned media experiments

Strains were inoculated in 15 mL ABB according to the standard overnight culture inoculation protocol. To prepare the conditioned media, 10 mL of the source organism cultures were transferred to 50 mL Falcon tubes and filtered in the anaerobic chamber using Steri-Flip (EMD Millipore). 10 mL ABB media was filtered as a control. Following pH measurements of the conditioned and unconditioned medias, 5 mL was filtered a second time. The remaining 5 mL of each conditioned media was adjusted to the pH of the filtered ABB media using 1M NaOH or 1M HCl and sterile filtered to represent the pH adjusted condition. The response organisms were normalized to an OD600 of 0.04 in ABB media. A Biomek 3000 liquid handling robot was used to transfer 60 μL of conditioned media into a 384-well plate (Corning) and 20 μL of the response organisms were added to a final OD600 of 0.01 in 75% conditioned media by volume. Plates were sealed (Diversified Biotech) and monitored every 30 min for 72 hr in a F200 Tecan Infinite Pro plate reader at 37°C.

### Bacterial culturing for metabolomics

A large batch of ABB media was prepared and used for all steps in the metabolomics experiments. ER and FP were grown in ABB media supplemented with 33 mM acetate (Sigma). Overnight cultures were grown for 72 hours at 37°C in 30 mL of ABB to saturation, diluted to 0. 01 OD600 and aliquoted into three 30 mL replicates in a 4-well plate (E&K Scientific). A well containing media was used to evaluate metabolite degradation as a function of time. Samples were collected an initial time point immediately following cell inoculation.

At each time point, the wells were mixed via serological pipet prior to sample collection and 800μL was removed for LC-MS. The samples were centrifuged at 6000 RCF for 5 min and 600 μL of supernatant was removed and filtered with a 0.22 μm filter unit (EMD Millipore). The supernatant was dispensed and simultaneously filtered into a microcentrifuge tube prior to freezing at -80°C. To elucidate metabolite profiles, samples were collected at varied time intervals for each organism prior to 24 hr except DP due to insufficient biomass accumulation for metabolomics measurements within the 24 hr time interval.

### Exo-metabolomics measurements

Media samples (0.5 mL) were lyophilized until dry in a Labconco 6.5 L lyophilizer (Labconco, Kansas City, MO) and stored at -80°C until extraction. In preparation for LC-MS analysis, the dried samples were resuspended in 200 μL MeOH containing internal standards (25 μM 3,6-dihydroxy-4-methylpyridazine, 4-(3,3-dimethyl-ureido)benzoic acid, d5-Benzoic acid, 9-anthracene carboxylic acid, ^13^C-glucose, ^13^C-^15^N-phenylalanine), bath sonicated for 20 mins, then centrifuge-filtered through a 0.22 μm PVDF membrane (Pall) and placed into glass HPLC vials.

Liquid chromatography tandem mass spectrometry (LC-MS/MS) was performed on extracts using an Agilent 1290 Ultra High Performance Liquid Chromatography stack (Agilent Technologies), with MS and MS/MS data collected using an Agilent 6550 Q-TOF (Agilent Technologies). Normal phase chromatographic separation was performed using a 150 × 2.1 mm, 5 μm, 200 Å SeQuant ZIC-pHILIC column (EMD Millipore) and guard column: 20 × 2.1 mm, 5 μm (EMD Millipore). The column was maintained at 40°C and solvent flow rate at a constant of 0.25 ml/min with a 2 μL injection volume for each sample. The HILIC column was equilibrated with 100% buffer B (90:10 ACN:H_2_O *w*/ 5 mM ammonium acetate) for 1.5 minutes, diluting buffer B down to 50% with buffer A (H_2_O *w*/ 5 mM ammonium acetate) for 23.5 minutes, down to 40% B over 3.2 minutes, to 0% B over 6.8 minutes, and followed by isocratic elution in 100% buffer A for 3 minutes. Samples were maintained at 4°C. Full MS spectra were collected from m/z 70-1050, with MS/MS fragmentation data acquired using 10, 20 and 40V collision energies at approximately 10,000 resolution. Metabolites were identified based on exact mass and retention time coupled with comparing MS/MS fragmentation spectra to purchased standards.

LC-MS data was analyzed using the Agilent MassHunter Qualitative Analysis (Agilent Technologies) followed by the Metabolite Atlas workflow^64^. A set of criteria was used to evaluate each of the detected peaks and assign a level of confidence in the compound identification. Compounds given a positive identification had matching retention time and m/z to a pure standard run using the same methods described above. A compound with the highest level of positive identification additionally had a matching MS/MS fragmentation spectrum to either an outside database (METLIN) or collected in house. Putative identifications were assigned to compounds with matching m/z and MS/MS spectrum.

### Model

The generalized Lotka-Volterra (gLV) model was used to represent microbial community dynamics. The gLV model is a set of coupled ordinary differential equations that represent the temporal variation in species abundance (*x_i_*). The model equations are:

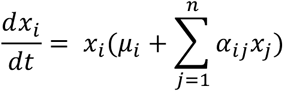

where *n*, μ, *α_ii_*, *α_ij_* represent the number of species, growth rates, intra-species and inter-species interaction coefficients, respectively. This model requires that the intra-species interaction coefficients are negative (*α_ii_* < 0). Inter-species interaction coefficients *α_ij_* can be positive or negative, representing a stimulatory or antagonistic microbial interaction.

### Parameter estimation and validation

Custom scripts in MATLAB (Mathworks) were used for model analysis and parameter estimation. A generalizable parameter estimation framework was developed to infer the 156 (*n*^2^ + *n*) parameters of the gLV model from time-series species abundance data. The experimental data included time-series measurements of OD600 for monospecies and communities and relative abundance of each species in the communities based on 16S rRNA gene sequencing. It is challenging to infer model parameters from compositional data generated by next-generation sequencing since this is an underdetermined problem and there are many solutions that would yield the same relative abundance output^59^. The absolute abundance of each species was estimated by the product of the relative abundance and the OD600 value (total biomass) at each time point. The nonlinear programming solver requires an initial point to the optimization problem. The initial point was computed by transforming the gLV system of equations into a linear system of equations:

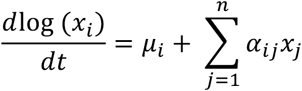

A linear least-squares algorithm (MATLAB) with bounds was used to solve for the unknown parameters using monospecies and time-series pairwise community measurements.

To minimize overfitting of the data, L1 regularization was used to penalize non-zero parameter values. Experiments were weighted equally in the objective function. A nonlinear programming solver (MATLAB) was used to minimize the following objective function:

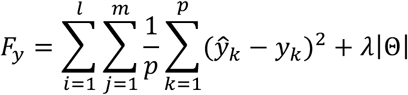

*l*, *m* and *p* denote community datasets, species in each community and time points, respectively. *λ* and Θ represent the regularization coefficient and the parameter vector. 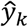 and *y_k_* designate the model prediction and experimental measurement of absolute abundance of a species at a specific time interval *k*.

In the optimization problem, an optimal *λ* value was identified to minimize overfitting of the data by balancing the goodness of fit and sparsity of the model. To do so, values of *λ* were scanned from 10^-5^ to 10. An optimization method was implemented to infer the best estimate of the model parameters using a specific value of *λ* (*λ* = 0.0077). The goodness of fit of the model was computed using the following equation:

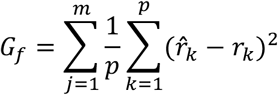

where *m*, *p*, 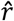 and *r* represent species, time points, predicted relative abundance and measured relative abundance, respectively. Validation of the model was performed using *G_f_* and the Pearson correlation coefficient between the model prediction and experimental data of all species in each community at each time point.

### Parameter uncertainty analysis

Bayes rule introduces the notion of prior and posterior information:

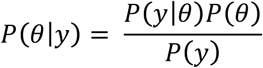

The parameters and data are represented by *θ* and y. We assume that the error is normally distributed with a mean of 0 using the following equation *y* = *𝑓*(*θ*) + *∈* where *∈*~*N*(0, *σ*^2^). The Posterior distribution *P*(*θ*|*y*) represents the uncertainty in the parameters. Since direct sampling from the Posterior distribution is challenging, the Metropolis-Hastings algorithm was used to estimate this distribution. Four independent chains were simulated for 500000 iterations from randomly sampled parameter values using a normal distribution with the mean equal to the best parameter estimate and standard deviation of 0.3 times the value of each parameter. A chain simulated for 500000 iterations from the single best estimate was also performed. A burn-in period of the first 300000 was excluded from the analyses to allow the chain to converge to the stationary distribution. The Gelman-Rubin potential scale reduction factor (PSRF) was used to evaluate convergence of the posterior distribution estimate. If the chains have converged to the target posterior distribution the PRSF should be close to 1. Our results showed that 81% of parameters had a PRSF less than 3.

## ACKNOWLEDGEMENTS

We would like to thank Nicholas Justice, Michael Fischbach and Justin Sonnenburg for helpful discussions. We are grateful to Ryan Clark, James Papadopolous, Joshua Hamilton and Susan Hromada for critical reading of the manuscript. This work was supported by the Defense Advanced Research Projects Agency (DARPA) Grant HR0011516183. O.S.V. was supported by the Simons Foundation at the Life Science Research Foundation postdoctoral fellowship.

## CONTRIBUTIONS

O.S.V. and A.P.A. designed the research. A.C.C., G.F. and R.H.H. carried out the experiments. O.S.V. designed and implemented computational modeling methods. O.S.V., A.C.C., R.H.H. and G.F. analyzed the data. O.S.V. wrote the manuscript and A.C.C., G.F., R.H. and A.P.A. assisted in revising the manuscript. T.N., O.S.V., A.P.A. and B.B. designed metabolomics experiments, R.L. performed LC-MS measurements and R.L. and B.B. assisted in analyzing the data.

## REFERENCES

1. Lozupone, C. A., Stombaugh, J. I., Gordon, J. I., Jansson, J. K. & Knight, R. Diversity, stability and resilience of the human gut microbiota. Nature 489, 220–230 (2012).

2. Tropini, C., Earle, K. A., Huang, K. C. & Sonnenburg, J. L. The gut microbiome: connecting spatial organization to function. Cell Host Microbe 21, 433–442 (2017).

3. Earle, K. A. et al. Quantitative Imaging of Gut Microbiota Spatial Organization. Cell Host Microbe 18, 478–488 (2015).

4. Louis, P., Hold, G. L. & Flint, H. J. The gut microbiota, bacterial metabolites and colorectal cancer. Nat. Rev. Microbiol. 12, 661–672 (2014).

5. Rooks, M. G. & Garrett, W. S. Gut microbiota, metabolites and host immunity. Nat. Rev. Immunol. 16, 341–52 (2016).

6. Ley, R. E. et al. Obesity alters gut microbial ecology. Proc. Natl. Acad. Sci. 102, 11070–5 (2005).

7. Sharon, G. et al. Specialized metabolites from the microbiome in health and disease. Cell Metab. 20, 719–730 (2014).

8. Fischbach, M. A. & Sonnenburg, J. L. Eating for two: how metabolism establishes interspecies interactions in the gut. Cell Host Microbe 10, 336–47 (2011).

9. Foster, J. a. & McVey Neufeld, K. A. Gut-brain axis: How the microbiome influences anxiety and depression. Trends Neurosci. 36, 305–312 (2013).

10. Donaldson, G. P., Lee, S. M. & Mazmanian, S. K. Gut biogeography of the bacterial microbiota. Nat. Rev. Microbiol. 14, 20–32 (2015).

11. Mark Welch, J. L., Hasegawa, Y., McNulty, N. P., Gordon, J. I. & Borisy, G. G. Spatial organization of a model 15-member human gut microbiota established in gnotobiotic mice. Proc. Natl. Acad. Sci. 201711596 (2017). doi:10.1073/pnas.1711596114

12. Sommer, F., Anderson, J. M., Bharti, R., Raes, J. & Rosenstiel, P. The resilience of the intestinal microbiota influences health and disease. Nat. Rev. Microbiol. 15, 630–638 (2017).

13. Ley, R. E., Peterson, D. a & Gordon, J. I. Ecological and evolutionary forces shaping microbial diversity in the human intestine. Cell 124, 837–48 (2006).

14. Faith, J. J. et al. The long-term stability of the human gut microbiota. Science (80-.). 341, (2013).

15. Relman, D. A. The human microbiome: Ecosystem resilience and health. Nutr. Rev. 70, S2–S9 (2012).

16. Faust, K. et al. Microbial co-occurrence relationships in the Human Microbiome. PLoS Comput. Biol. 8, (2012).

17. Mariat, D. et al. The Firmicutes/Bacteroidetes ratio of the human microbiota changes with age. BMC Microbiol. 9, 123 (2009).

18. Turnbaugh, P. J. et al. An obesity-associated gut microbiome with increased capacity for energy harvest. Nature 444, 1027–1031 (2006).

19. Foster, K. R. & Bell, T. Competition, not cooperation, dominates interactions among culturable microbial species. Curr. Biol. 22, 1845–50 (2012).

20. Hibbing, M. E., Fuqua, C., Parsek, M. R. & Peterson, S. B. Bacterial competition: surviving and thriving in the microbial jungle. Nat. Rev. Microbiol. 8, 15–25 (2010).

21. Bairey, E., Kelsic, E. D. & Kishony, R. High-order species interactions shape ecosystem diversity. Nat. Publ. Gr. 7, 1–7 (2016).

22. Gibson, T. E., Bashan, A., Cao, H. T., Weiss, S. T. & Liu, Y. Y. On the origins and control of community types in the human microbiome. PLoS Comput. Biol. 12, 1–21 (2016).

23. Faust, K. & Raes, J. Microbial interactions: from networks to models. Nat. Rev. Microbiol. 10, 538–550 (2012).

24. Astrom, K. J. & Murray, R. M. Feedback Systems: An Introduction for Scientists and Engineers. (Princeton University Press, 2010).

25. Qin, J. et al. A human gut microbial gene catalogue established by metagenomic sequencing. Nature 464, 59–65 (2010).

26. Scher, J. U. et al. Expansion of intestinal Prevotella copri correlates with enhanced susceptibility to arthritis. Elife 2, e01202 (2013).

27. Watterlot, L. et al. Faecalibacterium prausnitzii is an anti-inflammatory commensal bacterium identified by gut microbiota analysis of Crohn’s disease patients. Proc. Natl. Acad. Sci. 105, 16731–16736 (2008).

28. Fujimoto, T. et al. Decreased abundance of Faecalibacterium prausnitzii in the gut microbiota of Crohn’s disease. J. Gastroenterol. Hepatol. 28, 613–9 (2013).

29. Thota, V. R., Dacha, S. & Natarajan, A. Eggerthella lenta bacteremia in a Crohn’s disease patient after ileocecal resection. Future Microbiol. 6, 595–597 (2011).

30. Haiser, H. J. et al. Predicting and Manipulating Cardiac Drug Inactivation by the Human Gut Bacterium Eggerthella lenta. Science (80-.). 341, 295–298 (2013).

31. Larsen, N. et al. Gut microbiota in human adults with type 2 diabetes differs from non-diabetic adults. PLoS One 5, (2010).

32. Bucci, V. et al. MDSINE: Microbial Dynamical Systems INference Engine for microbiome time-series analyses. Genome Biol. 17, 121 (2016).

33. Widder, S. et al. Challenges in microbial ecology: building predictive understanding of community function and dynamics. ISME J. 10, 2557–2568 (2016).

34. de Vos, M. G. J., Zagorski, M., McNally, A. & Bollenbach, T. Interaction networks, ecological stability, and collective antibiotic tolerance in polymicrobial infections. Proc. Natl. Acad. Sci. 114, 201713372 (2017).

35. Gómez, J. M., Verdú, M. & Perfectti, F. Ecological interactions are evolutionarily conserved across the entire tree of life. Nature 465, 918–921 (2010).

36. Balzola, F., Bernstein, C., Ho, G. T. & Lees, C. A human gut microbial gene catalogue established by metagenomic sequencing: Commentary. Inflamm. Bowel Dis. Monit. 11, 28 (2010).

37. Murray, J. D. Mathematical Biology I: An Introduction. 17, (Interdisciplinary Applied Mathematics, 2002).

38. Vandeputte, D. et al. Stool consistency is strongly associated with gut microbiota richness and composition, enterotypes and bacterial growth rates. Gut 65, 57–62 (2016).

39. Roager, H. M. et al. Colonic transit time is related to bacterial metabolism and mucosal turnover in the gut. Nat. Microbiol. 1, 16093 (2016).

40. Bashan, A. et al. Universality of human microbial dynamics. Nature 534, 259–62 (2016).

41. Stein, R. R. et al. Ecological modeling from time-series inference: insight into dynamics and stability of intestinal microbiota. PLoS Comput. Biol. 9, e1003388 (2013).

42. Mounier, J. et al. Microbial interactions within a cheese microbial community. Appl. Environ. Microbiol. 74, 172–81 (2008).

43. Gábor, A., Villaverde, A. F. & Banga, J. R. Parameter identifiability analysis and visualization in large-scale kinetic models of biosystems. BMC Syst. Biol. 11, 54 (2017).

44. Rios-Covian, D., Gueimonde, M., Duncan, S. H., Flint, H. J. & De Los Reyes-Gavilan, C. G. Enhanced butyrate formation by cross-feeding between Faecalibacterium prausnitzii and Bifidobacterium adolescentis. FEMS Microbiol. Lett. 362, 1–7 (2015).

45. Duncan, S. H., Barcenilla, A., Stewart, C. S., Pryde, S. E. & Flint, H. J. Acetate utilization and butyryl coenzyme A (CoA): Acetate-CoA transferase in butyrate-producing bacteria from the human large intestine. Appl. Environ. Microbiol. 68, 5186–5190 (2002).

46. Wrzosek, L. et al. Bacteroides thetaiotaomicron and Faecalibacterium prausnitzii influence the production of mucus glycans and the development of goblet cells in the colonic epithelium of a gnotobiotic model rodent. BMC Biol. 11, (2013).

47. Filkins, L. M. et al. Coculture of Staphylococcus aureus with Pseudomonas aeruginosa drives S. aureus towards fermentative metabolism and reduced viability in a cystic fibrosis model. J. Bacteriol. 197, 2252–2264 (2015).

48. Flores, C. O., Poisot, T., Valverde, S. & Weitz, J. S. BiMat: A MATLAB package to facilitate the analysis of bipartite networks. Methods Ecol. Evol. 127–132 (2015). doi:10.1111/2041-210X.12458

49. Bucci, V. et al. MDSINE: Microbial Dynamical Systems INference Engine for microbiome time-series analyses. Genome Biol. 17, 121 (2016).

50. Friedman, J., Higgins, L. M. & Gore, J. Community structure follows simple assembly rules in microbial microcosms. Nat. Ecol. Evol. 1, (2017).

51. Momeni, B., Xie, L. & Shou, W. Lotka-Volterra pairwise modeling fails to capture diverse pairwise microbial interactions. Elife 6, 1–34 (2017).

52. Law, R. & Morton, R. D. Permanence and the assembly of ecological communities. Ecology 77, 762–775 (1996).

53. Mougi, A. & Kondoh, M. Diversity of Interaction Types and Ecological Community Stability. Science (80-.). 337, 349–351 (2012).

54. Coyte, K. Z., Schluter, J. & Foster, K. R. The ecology of the microbiome: Networks, competition, and stability. Science (80-.). 350, 663–666 (2015).

55. Faith, J. J., McNulty, N. P., Rey, F. E. & Gordon, J. I. Predicting a human gut microbiota’s response to diet in gnotobiotic mice. Science (80-.). 333, 101–4 (2011).

56. Arumugam, M. et al. Enterotypes of the human gut microbiome. Nature 473, 174–80 (2011).

57. Ley, R. E. Gut microbiota in 2015: Prevotella in the gut: choose carefully. Nat. Rev. Gastroenterol. Hepatol. 13, 69–70 (2016).

58. Kovatcheva-Datchary, P. et al. Dietary Fiber-Induced Improvement in Glucose Metabolism Is Associated with Increased Abundance of Prevotella. Cell Metab. 22, 971–982 (2015).

59. Fisher, C. K. & Mehta, P. Identifying keystone species in the human gut microbiome from metagenomic timeseries using sparse linear regression. PLoS One 9, 1–10 (2014).

60. Wright, E. S. & Vestigian, K. H. Inhibitory interactions promote frequent bistability among competing bacteria. Nat. Commun. 7, 1–7 (2016).

61. Guo, X. & Boedicker, J. Q. The contribution of high-order metabolic interactions to the global activity of a four-species microbial community. PLoS Comput. Biol. 12, 1–13 (2016).

62. Fadrosh, D. W. et al. An improved dual-indexing approach for multiplexed 16S rRNA gene sequencing on the Illumina MiSeq platform. Microbiome 2, 1–7 (2014).

63. Zhang, J., Kobert, K., Flouri, T. & Stamatakis, A. PEAR: A fast and accurate Illumina Paired-End reAd mergeR. Bioinformatics 30, 614–620 (2014).

64. Yao, Y. et al. Analysis of metabolomics datasets with high-performance computing and metabolite atlases. Metabolites 5, 431–442 (2015).

